# An Integrative Biology Approach to Quantify the Biodistribution of Azidohomoalanine *In Vivo*

**DOI:** 10.1101/2021.06.14.448308

**Authors:** Aya M. Saleh, Tyler VanDyk, Kathryn R. Jacobson, Sarah Calve, Tamara L. Kinzer-Ursem

**Affiliations:** Weldon School of Biomedical Engineering, Purdue University, 206 S. Martin Jischke Dr, West Lafayette, IN 47906; Purdue University Interdisciplinary Life Science Program, 155 S. Grant Street, West Lafayette, IN 47907; Paul M. Rady Department of Mechanical Engineering, University of Colorado – Boulder, 1111 Engineering Center, 427 UCB, Boulder, CO 80309

**Keywords:** Non-canonical amino acids, protein labeling, kinetics, compartment modeling, metabolomics

## Abstract

Identification and quantitation of newly synthesized proteins (NSPs) are critical to understanding protein dynamics in development and disease. Probing the nascent proteome can be achieved using non-canonical amino acids (ncAAs) to selectively label the NSPs utilizing endogenous translation machinery, which can then be quantitated with mass spectrometry. Since its conception, ncAA labeling has been applied to study many in vitro systems and more recently the in vivo proteomes of complex organisms such as rodents. In vivo labeling is typically achieved by introducing ncAAs into diet, which requires extended labeling times. We have previously demonstrated that labeling the murine proteome is feasible via injection of azidohomoalanine (Aha), a ncAA and methionine (Met) analog, without the need for Met depletion. With the ability to isolate NSPs without applying stress from dietary changes, Aha labeling can address biological questions wherein temporal protein dynamics are significant. However, accessing this temporal resolution requires a more complete understanding of Aha distribution kinetics in tissues. Furthermore, studies of physiological effects of ncAA administration have been limited to gross observation of animal appearance. To address these gaps, we created a deterministic, compartmental model of the biokinetic transport and incorporation of Aha in mice. Parameters were informed from literature and experimentally. Model results demonstrate the ability to predict Aha distribution and labeling under a variety of dosing paradigms and confirms the use of the model as a tool for design of future studies. To establish the suitability of the method for in vivo studies, we investigated the impact of Aha administration on normal physiology by analyzing the plasma metabolome following Aha injection. We show that Aha administration does not significantly perturb cellular functions as reflected by an unchanged plasma metabolome compared to non-injected controls.

**Author Summary:** As the machinery of life, proteins play a key role in dynamic processes within an organism. As such, the response of the proteome to perturbation is increasingly becoming a critical component of biological and medical studies. Dysregulation of protein mechanisms following exposure to experimental treatment conditions can implicate physiological mechanisms of health and disease, elucidate toxin/drug response, and highlight potential targets for novel therapies. Traditionally, these questions have been probed by studying perturbations in total proteins following an experimental treatment. However, the proteome is expansive and noisy, often an early response can be indiscernible against the background of unperturbed proteins. Here, we apply a technique to selectively label newly synthesized proteins, which enables capturing early changes in protein behavior. We utilize an amino acid analog that naturally incorporates into proteins, and investigate the tissue distribution, protein labeling efficiency, and potential physiological impact of this analog in mice. Our results demonstrate that we can reproducibly predict protein labeling and that the administration of this analog does not significantly alter in vivo physiology over the course of our experimental study. We further present a computational model that can be used to guide future experiments utilizing this technique to study proteomic responses to stimuli.

## Introduction

The use of non-canonical amino acid (ncAA) labeling for selective identification of newly synthesized proteins (NSPs) in mammalian cells was first introduced by Dieterich et al. in 2006 (1) and has since been applied to study several biological systems (see reviews (2, 3)). In this technique, an ncAA, typically a methionine (Met) analog, is introduced to the biological system of interest and incorporated into newly synthesized polypeptide chains using endogenous or engineered cellular translational machinery Distinction of nascent proteins from the constituent proteome is enabled by reactive chemical groups, such as azides and alkynes, which can be covalently modified via azide-alkyne cycloaddition (a click chemistry reaction) (2). As such, ncAA-labeled NSPs can be selectively conjugated to affinity or fluorescent tags for identification or visualization, respectively (1, 4). This technique has been successfully employed to probe protein dynamics in a variety of bacterial (5–7) and mammalian cells *in vitro* (8–10), as well as model organisms *in vivo*, including zebrafish (11) and *Xenopus* (12). More recently, ncAA labeling has also been shown to be effective in identifying NSPs in rodents (13–15). The expanding applications of ncAA labeling will enable previously inaccessible biological questions, wherein understanding the temporal dynamics of protein synthesis and turnover is critical, to be addressed.

For rodent proteome labeling, dietary administration of ncAA, typically enhanced with a Met-free diet, has been shown to achieve adequate labeling efficiency (13, 16). However, Met deprivation may affect normal physiology, particularly over longer labeling periods. Notably, the presence of Met in mammalian diet is essential for normal embryonic development (17–19), which constrains this method to studies of adult animals. In this regard, our group has previously demonstrated that labeling the adult and embryonic murine proteome can instead be achieved via systemic injection of ncAAs without the need for Met depletion (14, 20). Compared to feeding with an ncAA-enriched diet, the injection method achieves global proteome labeling in a shorter period of time, which enables the detection of proteins synthesized shortly after injection and proteins with high turnover rates (20). In addition, injections allow for accurate dosing calculations, which eliminates the inherent variability of the feeding method due to fluctuations in feeding patterns and intestinal absorption.

Despite the application of ncAA labeling in a number of studies to decipher complex cellular processes in animal models (13, 15, 16, 21), understanding of the kinetics of ncAA distribution in tissues, especially as it pertains to rates of protein incorporation and loss by degradation, is lacking. Determination of the timescale of ncAA uptake by tissues following administration and the lag time before maximum protein labeling are critical information for the design of robust temporal experiments to study the nascent proteome. Predicting ncAA pharmacokinetics in murine models will also enable optimizing the dosing regimen to attain the ideal concentrations to achieve sufficient protein labeling in the desired tissue over the course of the study.

In addition to the lack of knowledge of ncAAs distribution kinetics *in vivo*, evaluation of the physiological impact of ncAA administration to animals has been limited to examining changes in gross behavior, physical appearance and body weight (13–15). A more robust analysis of the effect of ncAA administration on the metabolome, and the corresponding implications for cellular function, is required to confirm the suitability of the method for *in vivo* studies.

The aim of this study was to characterize the distribution kinetics of azidohomoalanine (Aha), a widely used Met analog, in mice following subcutaneous injection, and to investigate the impact on normal physiology. To study the biodistribution of Aha, we measured the concentration in the plasma, liver, kidney, brain and skeletal muscle using liquid chromatography-tandem mass spectrometry (LC-MS/MS) over a period of 24 h. This dataset was used to develop a deterministic compartment model of small molecule biokinetics that characterizes the movement of freely diffusive Aha (fAha) throughout the mouse circulatory system and into tissues. In addition, we used fluorescent western blotting to measure protein labeling in these tissues during the same period of time. This second dataset was used to inform a model of relative protein labeling as a function of fAha availability to characterize both the predicted labeling profile for a given experimental treatment of Aha and the relative synthesis and turnover rates of Aha-labeled proteins. We demonstrate that this model can be used to characterize nascent protein synthesis and turnover within distinct tissues. Furthermore, we validated the capability of this model to predict NSP labeling under more complex, multiple injection dosing paradigms, which can be a tool to guide the design of future experiments utilizing Aha labeling.

We also probed the effect of Aha incorporation into NSPs and investigated whether Aha labeling perturbs normal physiological functions. We compared the plasma metabolome 24 h after Aha injection to that of non-injected mice to identify if metabolic pathways were dysregulated due to protein labeling with Aha. Only ~ 1.3% of metabolites were differentially regulated in the injected mice, indicating that Aha administration does not have a significant impact on normal physiology. Taken together, these results provide a fundamental understanding of the interrelation between the distribution kinetics of the ncAA into murine tissues and the associated degree of protein labeling, as well as the impact of ncAA injection on physiological functions.

## Results and Discussion

Effective protein labeling is critical to enrich Aha-labeled proteins with high signal-to-noise ratio for accurate quantitative MS measurements and identification of newly synthesized proteins. Depending on the tissue type and biological processes to be studied, multiple injections of Aha may be required to attain a high enough degree of labeling that results in a suitable MS signal. Therefore, optimizing the dose and frequency of Aha injections is critical for the appropriate design of labeling studies. In this regard, we sought to describe the distribution and labeling kinetics of Aha with an experimentally informed deterministic model. Our model was developed to describe: (1) the transport of freely diffusive Aha (fAha), Aha that is yet to be incorporated into protein, in the plasma, (2) the circulatory exchange of fAha into tissues of adult mice and (3) the degree of Aha incorporation into proteins (pAha) in specific tissues.

### Kinetic Model of Aha Biodistribution Describes Transport and Exchange

To capture the dynamics of fAha distribution *in vivo*, an *in silico* model system of ordinary differential equations (ODEs) was generated describing the physiological processes of small molecule transport. Dosed fAha was introduced into the model at a non-localized reservoir, to mimic the injection site of our subcutaneous dosing paradigm. From this reservoir, fAha enters the murine circulatory system, at a rate that is a function of reservoir concentration and transport kinetics, and is distributed to distinct tissue compartments (Figure 1).

**Figure 1.**
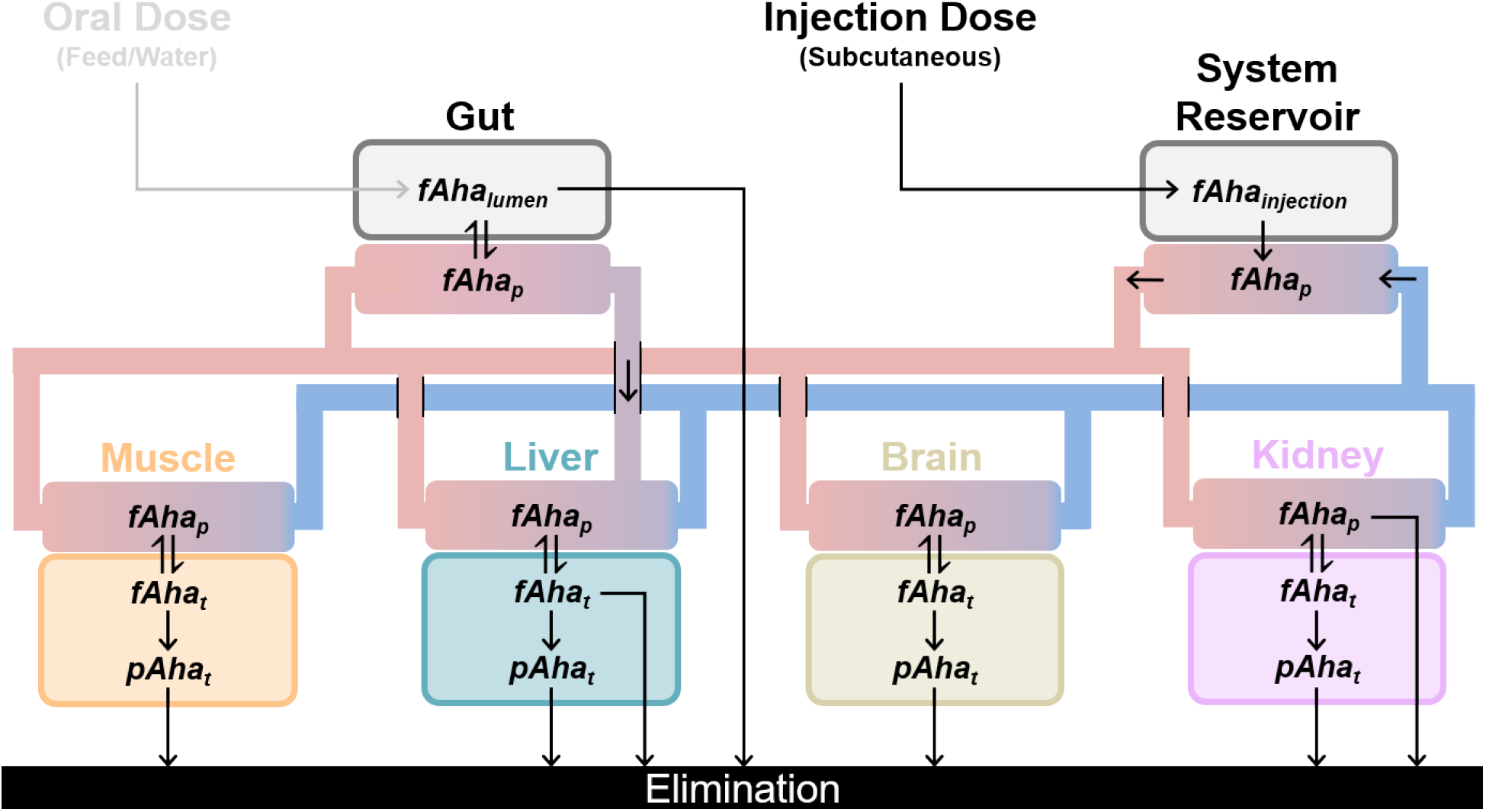
Biodistribution of fAha via transport and exchange. Introduced at a distinct injection site, fAha is allowed to enter the circulation at a systemic venous reservoir. Driven by circulation, fAha is passed through the arterial system (red) into distinct tissue compartments where exchange occurs at tissue specific rates. Arrows indicate directional movement of fAha. Each tissue compartment consists of two sub-compartments: plasma available for surface exchange (red to blue gradient) and an intracellular volume (illustrated here as the bottom compartments: from left to right muscle, liver, brain, and kidney). A mechanism for oral dosing (via plasma exchange with the gastrointestinal lumen) is illustrated, but not included in this model.

Within each compartment, the time-dependent rate of change of the fAha plasma concentration ([*fAha_p_*]) available for exchange with each tissue can be described as a mass balance with two stages: transport and exchange. The transport stage is governed by circulatory blood flow.

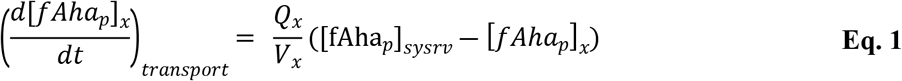

where *Q_x_* is the blood flow rate between tissue ‘*x’* and a systemic venous reservoir (*sysrv*), and *V_x_* is the corresponding volume of plasma relevant to each tissue (*Q/V* represented as a lumped constant *qb* in Supplements S2,S3). All kinetic parameters for circulatory transport were normalized by tissue mass to compare relative perfusion rates between tissue compartments of differing size. Once localized to a tissue, fAha in the plasma can also be exchanged across the cell membrane with the intracellular Aha concentration ([*fAha_t_*]) and is described as follows:

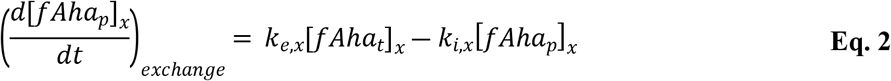

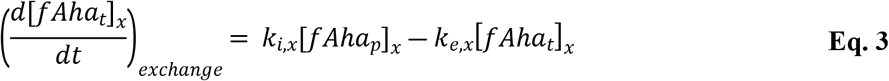

where *k_i,x_* and *k_e,x_* are the tissue specific import and export rates (min^−1^⋅mg^−1^) for fAha across the cell membrane. The liver tissue, gut and kidney plasma compartments were assigned additional system removal terms (*k_r,x_*, Supplements S2,S3) accounting for excretion and metabolization of fAha. The two stages of distribution were combined into a single system of ODEs, parameterized and bound within reasonable ranges for a model of small molecule pharmacokinetics (Supplements S2-S5) (22–25).

### Kinetic Model of Protein Labeling Captures Aha Incorporation

Within each tissue compartment, *fAha_t_* is incorporated into proteins via protein synthesis. As a Met analog, Aha is able to bind to methionyl-tRNA synthase, albeit at a much slower rate (*k_cat_·K_m_*^−1^ Aha: 1.42E-3, Met: 5.47E-1 μM^−1^·s^−1^) (26). Because the rate constant of Aha binding to the methionyl-tRNA synthase is much slower than Met, and previous estimates of the amount of Aha incorporation into NSPs was less than 10% (14), we made a modeling assumption that [*fAha_t_*] is negligibly depleted by incorporation into protein (further justification, Supplement S2) and that recycling of Aha is nonexistent.

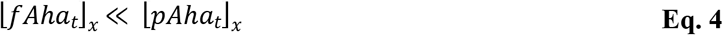

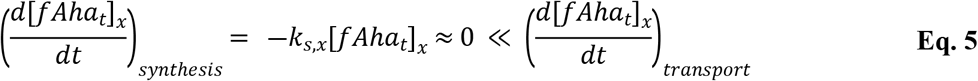

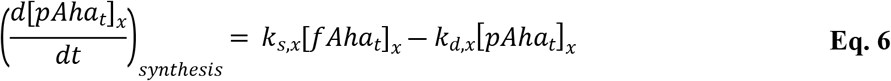

where *k_s,x_* and *k_d,x_* are the tissue-specific (denoted for tissue ‘*x*’ as above) rate constants of incorporation of Aha into proteins due to synthesis and loss of Aha-labeled proteins due to degradation. These equations describing Aha labeling of proteins in each tissue were added to the biodistribution model establishing a time resolved predictive model of tissue-specific protein labeling given a variety of input dosing paradigms.

### Experimental LC-MS/MS and Western Blotting Data Enables Parameter Fitting

Model parameters were initialized and bound within reasonable ranges, informed from literature and experimental measurements as described in the methods (Supplements S3-5) (22–25), then underwent least squares regression to match experimentally measured data of Aha concentration and labeling. To inform fitting, fAha concentration profiles in plasma and tissues were determined by injecting Aha subcutaneously into mice at 0.1 mg·g^−1^ total body weight and sacrificing 0.5-24 h post injection (hpi). Accurate identification of fAha in each tissue was performed using LC-MS/MS multiple reaction monitoring (Figure 2A). Additionally, the kinetics of Aha incorporation into tissue proteins were described by examining the degree of protein labeling within each tissue over the duration of the study. To this end, tissue homogenates were reacted with biotin-alkyne via copper-catalyzed click reaction, analyzed by western blotting using a fluorescent streptavidin conjugate and the change in fluorescence intensity relative to the non-injected controls was measured as an analog for pAha labeling (Figure 2B,C).

**Figure 2.**
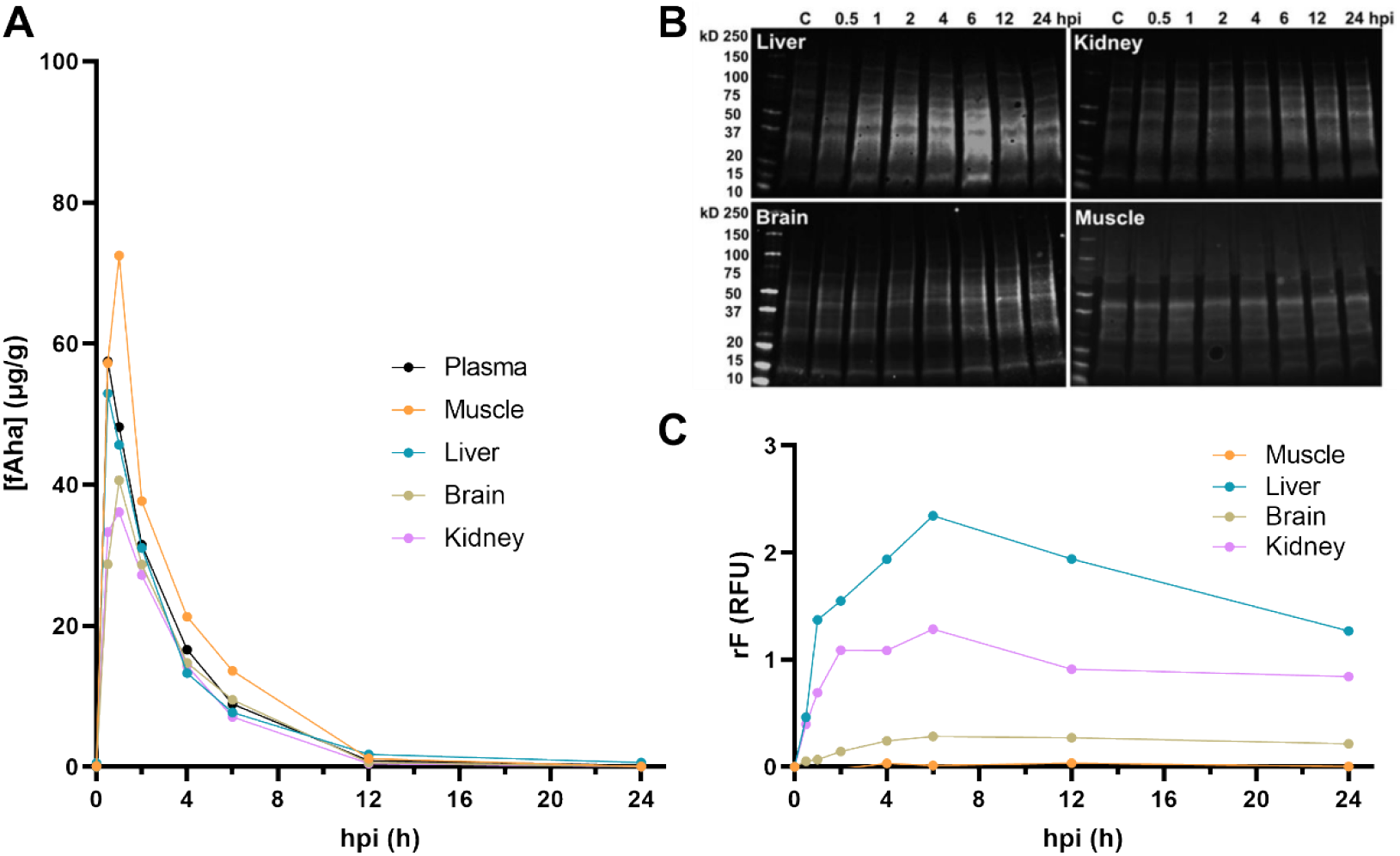
Free Aha concentration and protein incorporation and turnover kinetics in murine tissues were determined experimentally to inform the pharmacokinetics model. **(A)** The concentration profile of fAha in the plasma and different tissues. The amount of Aha (μg) measured by LC-MS/MS was normalized by the total plasma volume or tissue mass and averages were plotted over time. **(B)** Fluorescent western blots of the tissue homogenates of control non-injected samples (C) and samples collected 0.5 – 24 h post Aha injection (hpi). **(C)** Fluorescence intensity of western blot lanes were normalized to that of the respective control samples and averages were plotted as function of time (n=3 biological replicates).

The degree of fluorescent signal normalized relative to the background (*rF*) measured in relative fluorescence units (RFU) by semi-quantitative western blotting was assumed to be linearly proportional to the concentration of pAha using a fluorescent labeling factor, *k_f_*. For each tissue (denoted ‘*x*’)

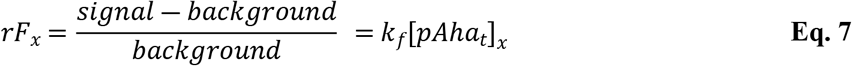

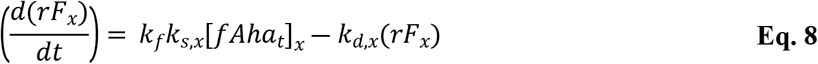

### Aha Labeling Captures Relative Protein Synthesis and Turnover Dynamics in Murine Tissues

The concentration profiles of fAha in the plasma and tissues peaked between 0.5 and 1 hpi and was mostly cleared from the system by 12 hpi, with the liver having the earliest peak compared to the other tissues (Figure 3A-D). The early peak can be attributed to the high blood perfusion of the liver (27), which likely results in faster distribution equilibrium of Aha into the liver compared to other tissues.

**Figure 3.**
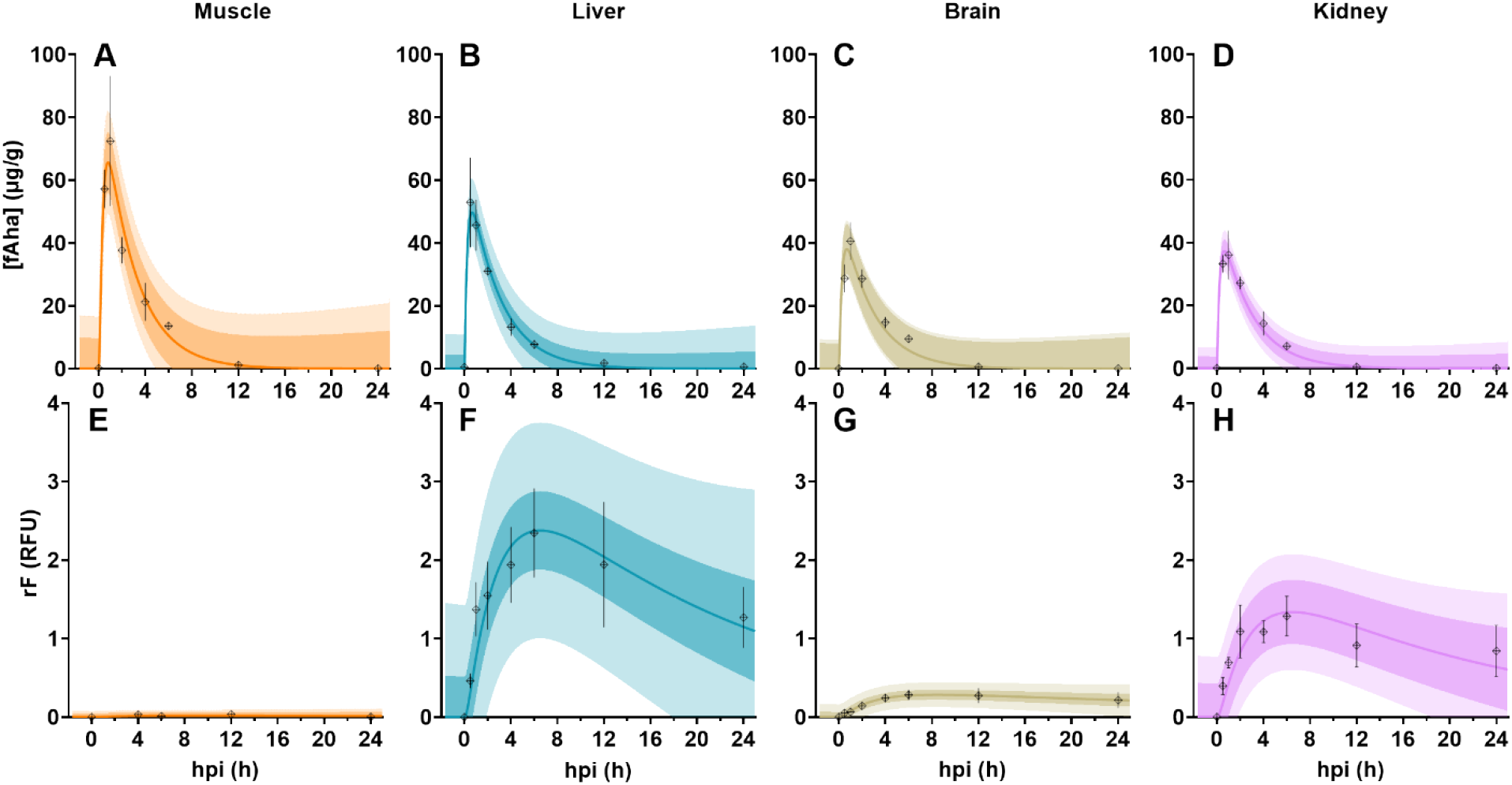
Aha kinetics and protein labeling following subcutaneous injection of Aha. **(A-D)** The concentration profiles of free Aha (fAha) in the plasma and different tissues. The amount of Aha (μg) measured by LC-MS/MS analysis was normalized by the total tissue mass and plotted over time. **(E-H)** Relative fluorescence (rF), as measured by western blotting, of proteins isolated from each tissue as a function of time. Filled points represent mean experimental measurement at each time point, error bars indicate experimental standard deviation (n = 3 biological replicates). Colored traces indicate best fit of model to each dataset, with darker and lighter shaded regions showing 95% prediction intervals for residual error calculated from mean experimental values and all experimental replicates, respectively.

In all tissues, maximum protein labeling was observed around 6 hpi (Figure 3E-H). However, the degree of labeling, represented by the maximum fold increase in fluorescence intensity compared to an internal control, and the kinetics of protein incorporation and turnover varied considerably between tissues. The liver showed the highest degree of labeling (Figure 3F) as well as the highest relative rates of Aha incorporation and protein turnover (Table 1), whereas skeletal muscle had the lowest degree of labeling (Figure 3E) and the slowest relative rates of incorporation and turnover (Table 1). Interestingly, the amount of fAha (μg) per unit mass tissue (g) was higher in skeletal muscle than in liver (Figure 3A,B), indicating that the low degree of labeling observed in skeletal muscle is predominantly due to a slow rate of muscle Aha incorporation, implying a lower rate of Met incorporation and protein synthesis.

**Table 1.** Protein incorporation and degradation estimated from Aha dynamics. Best fit parameter values for Aha incorporation, and protein turnover and half-lives in studied tissues.

| Parameter, Units | Muscle | Liver | Brain | Kidney |
| --- | --- | --- | --- | --- |
| Protein turnover rate, $h^{-1}$ | 2.23E-2 | 4.97E-2 | 2.12E-2 | 5.06E-2 |
| Protein half-life, h | 31.05 | 13.94 | 32.6 | 13.69 |
| Aha Incorporation*, RFU($\mu g$ pAha · h)$^{-1}$ | 5.36E-4 | 2.28E-2 | 6.06E-3 | 5.43E-2 |

\*Relative incorporation rate = $k_f \cdot k_s \cdot mt^{-1}$ , units include RFU per mass pAha.

The observed differences in fluorescence intensities are in agreement with previous isotope labeling studies that showed faster protein turnover rates in liver and kidney compared to brain and skeletal muscle (28–30). Notably, our model estimated a protein half-life in the brain that is 2.7 times higher than the liver (Table 1), in close agreement with the findings of Price et al. (9 h^−1^ and 3 h^−1^ days for brain and liver, respectively) (29). The discrepancies between previously reported values and the half-lives values estimated here can be attributed to the shorter timescale of our experimental setup as compared to stable isotope labeling technique and the use of semi-quantitative western blotting measurements rather than MS. A more robust quantitation of pAha concentrations using MS will be required for a more precise determination of absolute protein kinetics in different murine tissues.

### Descriptive Statistics Support Model of Biodistribution and Labeling

Model validity was examined using the following metrics to investigate parameter stability and goodness of fit. First, a prediction interval of residuals was generated for each tissue studied, for both the biodistribution and protein labeling models. A 95% prediction interval (PI_α=0.05_) of residuals was calculated using all experimental replicates (Figure 3). A second, tighter prediction interval was calculated using the mean for each time point (Figure 3). For each tissue, the width of PI_α=0.05_ from the average line of best fit (*ŷ*) can be approximated using a naively informed forecast interval that assumes a normal distribution of residual error (31).

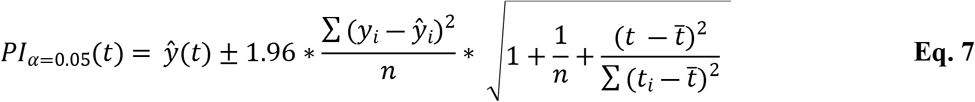

where, *n* is the total number of observations, (*t_i_, y_i_*) are the coordinates for each observation, *t* is the time point of the predicted residual, 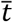 is the average time of all experimental observations. These prediction intervals demonstrate a narrow range of residual error and capture all experimental means and >95% of experimental values, indicating an accurate predictive model.

Second, the covariance matrix of least squares regression was used to inform a standard error of fitting (SE_f_) for all fitted parameters in the biodistribution model (Supplements S3-S5). SE_f_ describes variability of each parameter, but also reflects upon the definition of the model. Parameters with best fit values near to the constraints are less predictable; wide error would be indicative of a poorly constrained system. Error also increases with the number of fitted parameters, is inversely related to the quantity of data points available for fitting, and is weighted by the metric used for optimization. Among all fit parameters, we found relatively few SE_f_ values greater than best fit values by more than an order of magnitude (Figure 3, Supplements S3-5). Notably, we found that the widest error ranges in parameters that were the least well characterized in the literature, specifically rates describing tissue import and export of fAha. Low SE_f_ values, particularly among parameters within reported literature values, support a well characterized model, although this metric is not fully sufficient to describe a complex non-linear regression.

A more thorough analysis was achieved using Monte-Carlo Latin hypercube sampling (LHS) to perform efficient sampling of the input parameter space and correlation with partial rank correlation coefficients (PRCCs). This global sensitivity analysis was performed against several metrics to probe the influence of variation in parameter values on (1) model fitness (2) predicted Aha labeling levels in each tissue (rF). To address the first of these, PRCC values, which vary between 1 (perfect positive correlation) and −1 (perfect negative correlation), were calculated comparing variation in the input parameter values against the sum of square errors for each fitting model. This analysis generates a PRCC value for each parameter (PRCC_f_) that characterizes its relative effect on the fitting metric, and therefore the fitting process. PRCC_f_ values higher in absolute magnitude indicate parameters that, when varied independently, have the greatest influence on the fitting metric with values > 0.8 considered as indications of regions of instability in the model (32, 33). All parameters fall safely below a PRCC_f_ value of 0.3. Parameters with the highest PRCC_f_ values are those that influence the system removal of fAha, likely due to the rapid metabolic profile exhibited following subcutaneous injection (Figure 3). PRCC_f_ values for elimination and import rates into tissues that work actively to remove fAha were > 0.2.

Taken together, PRCC_f_ and SE_f_ values demonstrate that influential parameters are not those with the widest error ranges (Figure 4, Supplements S5-S7). Furthermore, when comparing these values across the parameter space, there is no one tissue that exhibits over-sensitive behavior by either metric, beyond systemic elimination. These findings provide support for the model definition, boundary constraints, and biological relevance of best fit parameters.

**Figure 4.**
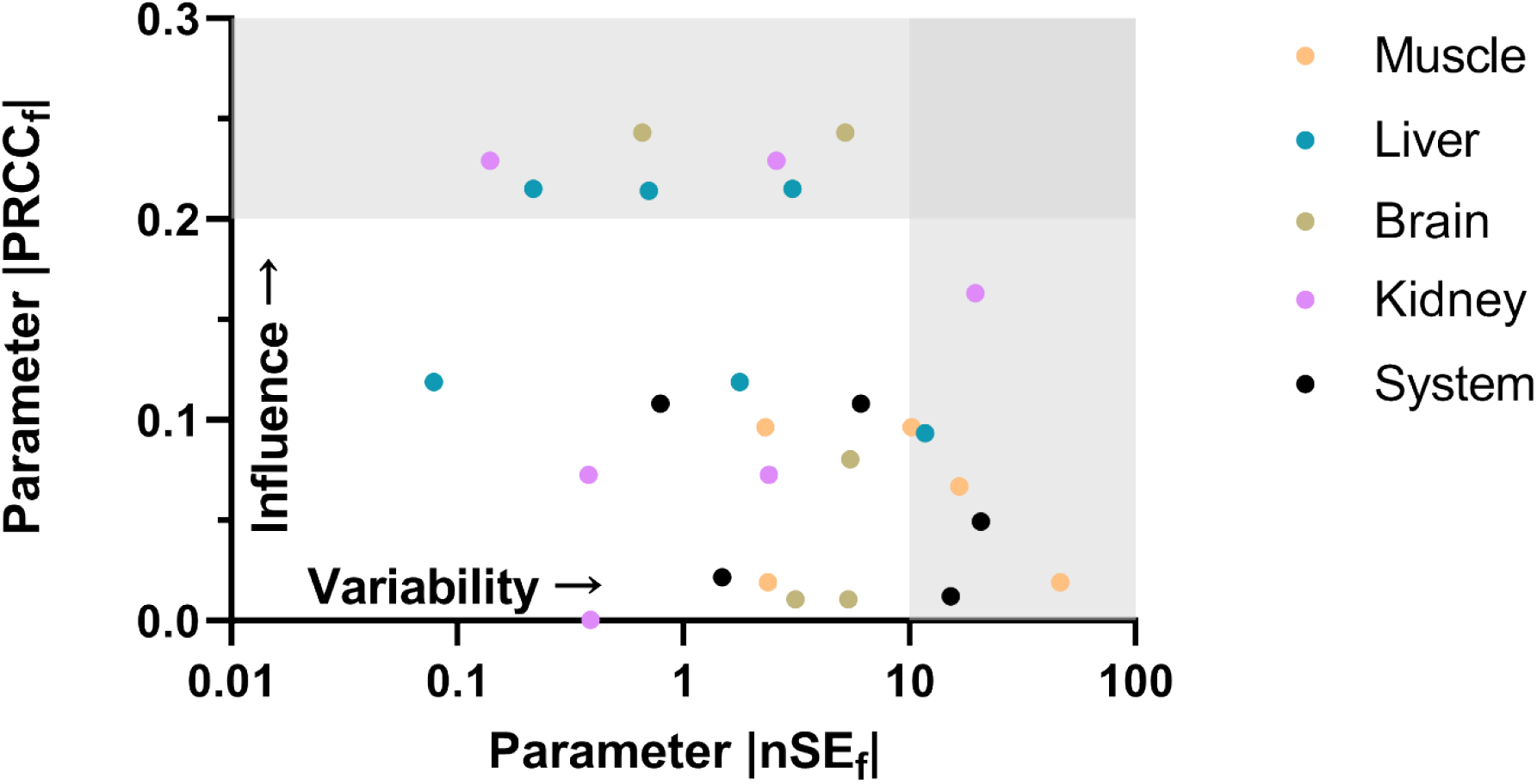
Sensitivity analysis for model parameter fitting. Each point represents a distinct parameter from the model (colored by associated tissues) and coordinates indicate absolute values of normalized standard error of fitting (nSE_f_ = SE_f_ / best fit value) on the x-axis and PRCC_f_ on the y-axis. Light gray shaded regions indicate parameters that have either a relatively high model influence (PRCC_f_ > 0.2) or variability (nSE_f_ > 10). The dark grey shaded region indicates a domain where parameters would be considered unstable or poorly constrained with both a high variability and influence on the model.

Further application of PRCC analysis allowed examination of the influence of parameter values upon the predicted degree of Aha labeling in each tissue over time. All model parameters were varied within their estimated fitting range (Supplements S5-S7) and sampled via LHS (n=10000). An rF value in each tissue was predicted for each timepoint from 0-24 hpi and the resulting output was correlated to the input parameter variation using PRCC analysis. Resulting traces elucidate kinetic parameters and physiological mechanisms that most influence rF during different time domains (Figure 5). For example, immediately after injection (0-4 hpi), the rate of absorption of Aha from initial injection site (*kaSubcu*) is a critical mechanism driving rapid labeling efficiency in all four studied tissues. However, the importance of this parameter declines over time, just as the influence of other parameters increases, particularly those representing Aha degradation and elimination. The influence of liver transport is prominent in all four tissues indicating the liver plays a major role in Aha kinetics. This suggests Aha may be primarily metabolized and eliminated by the liver, rather than by eliminated via excretion. This possibility is consistent with the known liver functions of toxin removal and amino acid metabolism (34).

**Figure 5.**
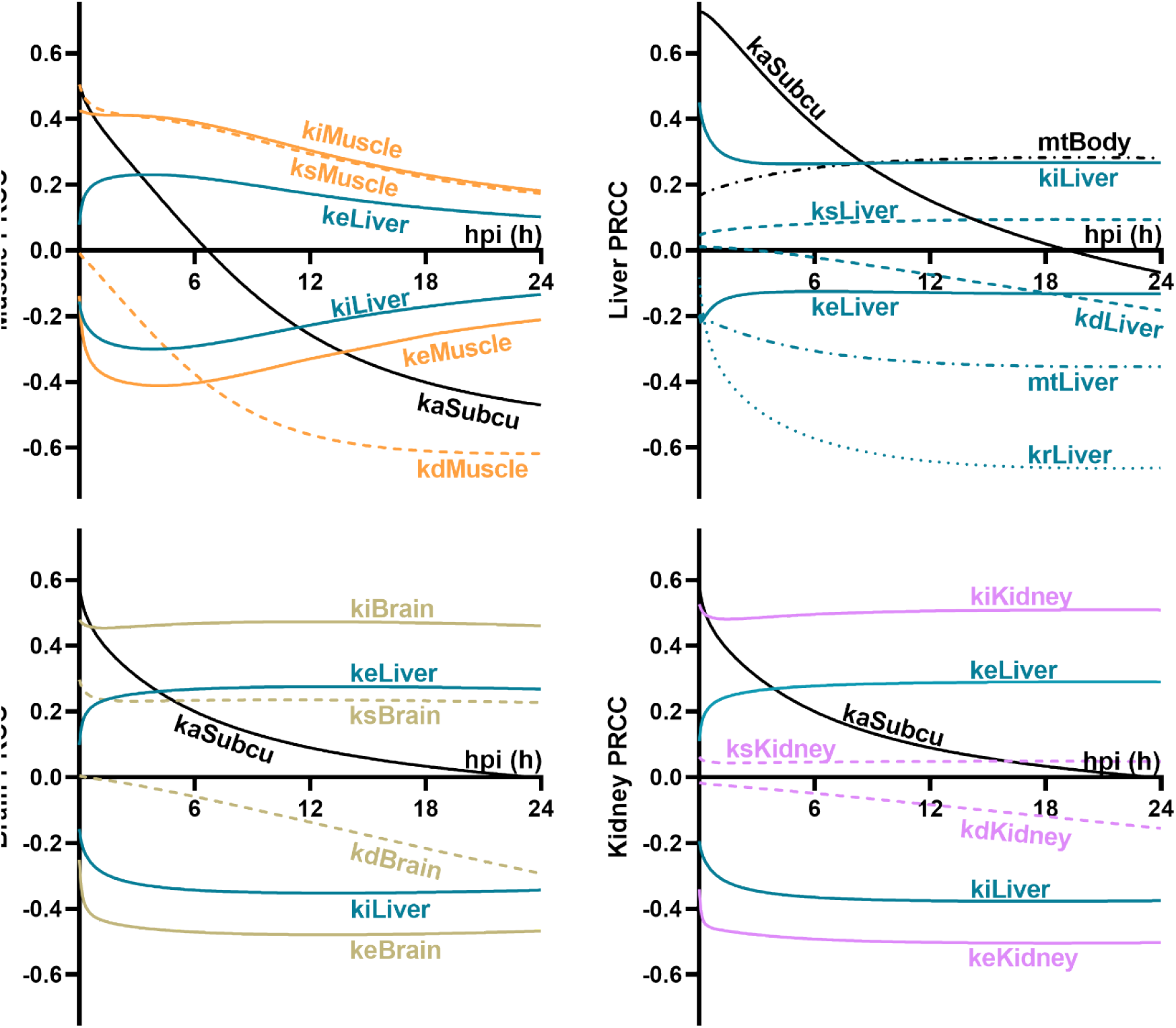
Sensitivity analysis of input parameter variation and Aha labeling in tissues over 24 hpi. Each plot shows the time resolved PRCC values for a selection of parameters using predicted rF in each tissue as a correlation metric. Parameters shown are either (1) descriptive of dynamics within the relevant tissue or (2) related to a different tissue, but with a significant influence on the tissue of interest based upon PRCC values > |0.2|. Trace colors indicate associated tissues (black = systemic param8883eter, blue = liver, orange = muscle, green = brain, purple = kidney), trace patterns indicate parameter type (solid = fAha transport, dotted = fAha elimination, dashed = pAha synthesis/degradation, dot-dashed = tissue mass).

### Predictive Simulations Accurately Capture Alternate Dosing Paradigms

Attaining sufficient protein labeling is critical for accurate identification and quantitation of Aha-labeled proteins using LC-MS/MS. For instance, note the near negligible labeling of skeletal muscle in Figure 3. If muscle labeling is desired, a much higher Aha dose might be required to attain sufficient labeling. This underlines the importance of tailoring the dosing regimen of Aha (*i.e.* amount per dose and dosing frequency) to the tissue of interest and the biological question under investigation. Using the model described above, fAha biodistribution and tissue protein labeling can be predicted for alternative dosing regimens to aid future experimental design and predict labeling efficiency depending on the conditions of a study. However, the model was based on the concentration of Aha in plasma and tissues over 24 h after a single subcutaneous injection. While this data was sufficiently robust to develop a well-informed model of Aha patterning after one subcutaneous dose, the use of this model to predict Aha content for longer time scales or multiple dose paradigms would generate naïve forecasts. To address this gap, we validated the predictions of our model against a data set that spans a longer time period and multiple injected doses. The model was used to predict the rF labeling of brain and liver tissues for two alternative dosing paradigms with either 12 h repeated doses (hrd) or 24 hrd over a 36 h period (Figure S7). As an internal control for western blotting variation, these new experimental values were normalized by a shared time point with the previous study (6 hours post initial injection) for each tissue (Figure 6).

**Figure 6.**
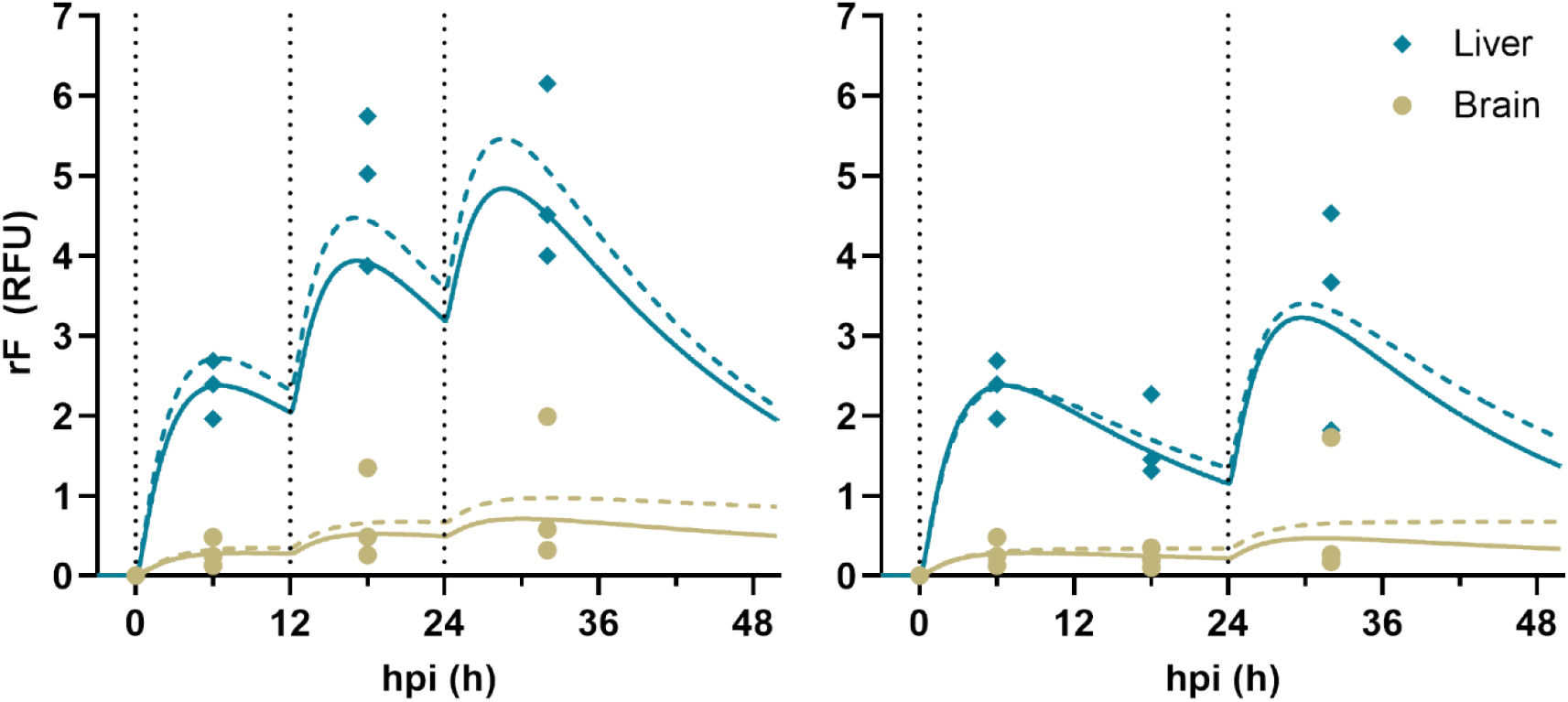
Model accurately predicts relative labeling in the brain and liver with multiple injection doses. rF experimental data (dots) and model predictions (lines) for **(A)** 12 hrd and **(B)** 24 hrd. For each tissue, experimental replicates from the repeated dose study are displayed at 6, 18, and 32 hpi as individual points. The solid line is the predicted trace using the best fit model from the original robust dataset. The dashed line is the model with parameter values refit to the data from the repeated dose experiments. The vertical dotted line indicates dose injection timepoints.

To determine the ability of the original model to predict Aha incorporation into proteins in various tissues, the data from each repeated dose study was used to refit the relative pAha synthesis rate and degradation for each tissue under each repeated dose paradigm. Relative to the parameter fit with the original experimental data, there was only a slight reduction in the standard error of regression (SE_reg_), a goodness-of-fit metric, between the original and refit parameters in each tissue (Table 2). Additionally, among all refit parameters, a single parameter was adjusted beyond a single standard error of fit (SE_f_) from the original best fit value (12 hrd liver Δ(*k_f_ ·k_s_*) ≈ +1.97SE), and only the degradation rate in the brain changed by >20% (Table 2).

**Table 2.**
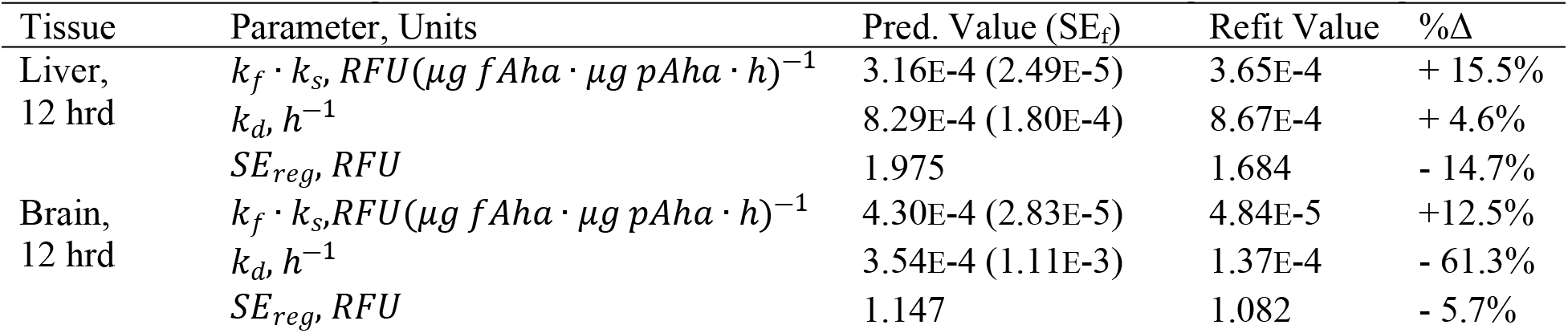
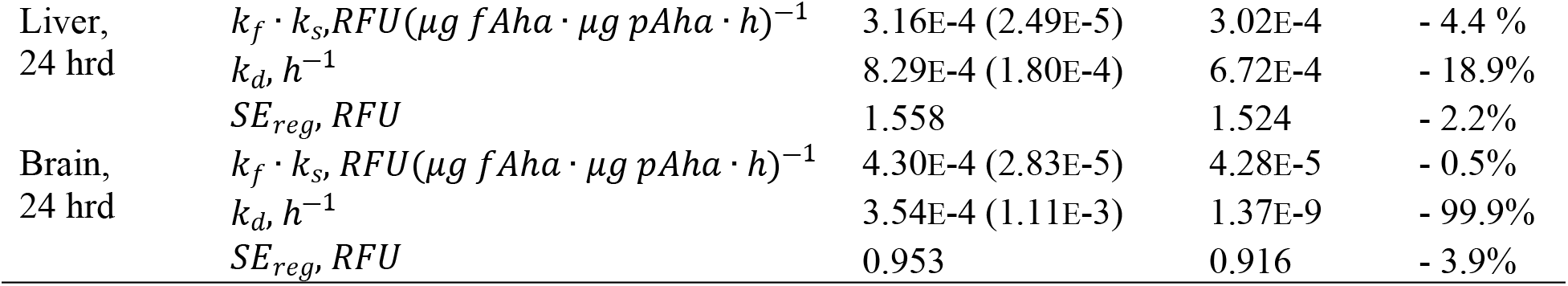
Parameter and goodness-of-fit statistics related to the alternative dosing models in Figure 6.

| Tissue | Parameter, Units | Pred. Value ($SE_f$) | Refit Value | %$\Delta$ |
| --- | --- | --- | --- | --- |
| Liver,<br>12 hrd | $k_f \cdot k_s, RFU(\mu g fAha \cdot \mu g pAha \cdot h)^{-1}$ | 3.16E-4 (2.49E-5) | 3.65E-4 | + 15.5% |
| | $k_d, h^{-1}$ | 8.29E-4 (1.80E-4) | 8.67E-4 | + 4.6% |
| | $SE_{reg}, RFU$ | 1.975 | 1.684 | - 14.7% |
| Brain,<br>12 hrd | $k_f \cdot k_s, RFU(\mu g fAha \cdot \mu g pAha \cdot h)^{-1}$ | 4.30E-4 (2.83E-5) | 4.84E-5 | +12.5% |
| | $k_d, h^{-1}$ | 3.54E-4 (1.11E-3) | 1.37E-4 | - 61.3% |
| | $SE_{reg}, RFU$ | 1.147 | 1.082 | - 5.7% |
| Liver,<br>24 hrd | $k_f \cdot k_s, RFU(\mu g fAha \cdot \mu g pAha \cdot h)^{-1}$ | 3.16E-4 (2.49E-5) | 3.02E-4 | - 4.4 % |
| | $k_d, h^{-1}$ | 8.29E-4 (1.80E-4) | 6.72E-4 | - 18.9% |
| | $SE_{reg}, RFU$ | 1.558 | 1.524 | - 2.2% |
| Brain,<br>24 hrd | $k_f \cdot k_s, RFU(\mu g fAha \cdot \mu g pAha \cdot h)^{-1}$ | 4.30E-4 (2.83E-5) | 4.28E-5 | - 0.5% |
| | $k_d, h^{-1}$ | 3.54E-4 (1.11E-3) | 1.37E-9 | - 99.9% |
| | $SE_{reg}, RFU$ | 0.953 | 0.916 | - 3.9% |

### Aha Administration Does Not Perturb Normal Physiology in Mice

In addition to the characterization of Aha distribution kinetics in mice, the impact of Aha administration on normal physiology must be qualified to establish the applicability of the method for *in vivo* studies. To this end, the physiological impact of Aha incorporation into newly synthesized proteins was evaluated using untargeted plasma metabolomic analysis. Since metabolites are the end products of cellular biological processes, we reasoned that a lag time is expected between potential changes in protein functions due to Aha incorporation and any associated effects on metabolism. Therefore, given that maximum protein labeling occurred ~ 6 hpi (Figure 2B,C and Figure 3E-H), we analyzed the plasma metabolome 24 hpi to identify any potential changes in metabolic pathways in response to Aha incorporation.

LC-MS metabolomic analysis of the plasma identified a total of 1268 mass features (*i.e.* metabolites). The peak area of each mass feature is proportional to the amount of the corresponding ion in the sample and was used as a measurement for the relative abundance of each identified metabolite across samples. Principal component analysis (PCA) revealed no distinct segregation between the control and Aha mice, indicating that there were no global differences in the plasma metabolome between the two groups (Figure 7A).

**Figure 7.**
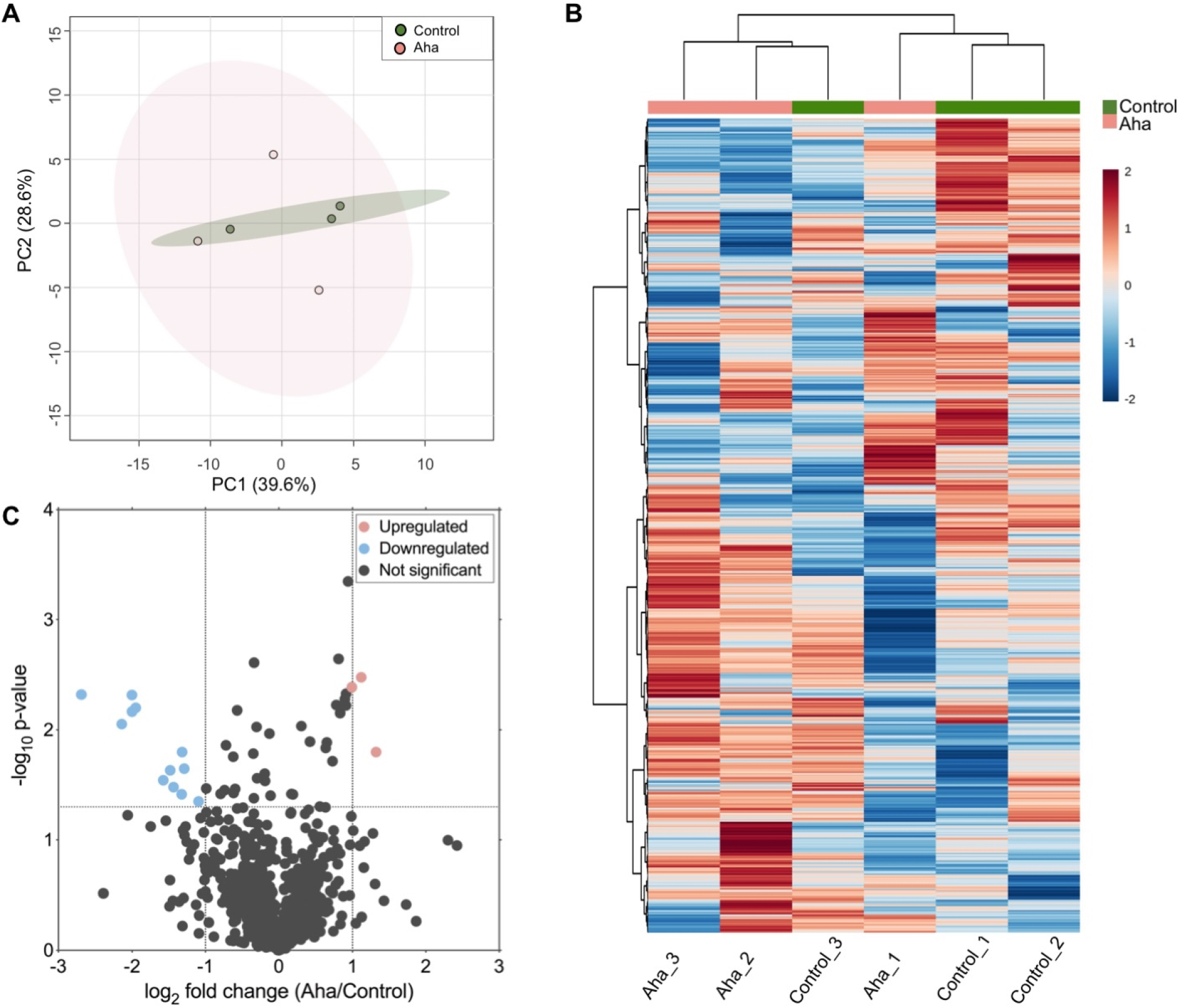
Aha administration does not significantly change the murine plasma metabolome. **(A)** Principal component analysis (PCA) showed no clear separation between control and Aha-treated groups in the first two components. Components 1 and 2 account for 39.6% and 28.6% of the total data variability, respectively. Green and red dots denote control and Aha samples, respectively. Green and red shaded areas represent the 95% confidence bands of the control and Aha samples, respectively. **(B)** Heat map of unsupervised hierarchal clustering analysis (HCA) of the identified metabolites show lack of clustering between replicates of each group. Color scale indicates metabolite abundance; blue: lowest, red: highest. **(C)** Volcano plot comparing the relative abundance of the identified metabolites between control and Aha groups according to statistical significance and fold change. Horizontal line indicates *p*-value = 0.05 and vertical lines indicate ± 2-fold change. Grey, red and blue circles denote equally-abundant, upregulated and downregulated metabolites, respectively.

It should be noted that PCA cannot be performed in the presence of missing values. The occurrence of missing values is common in untargeted metabolomic data, resulting from the presence of metabolites with concentrations that are lower than the MS detection limit or due to technical reasons such as incomplete ionization or inaccurate peak detection [35]. In our dataset, a total of 194 (2.5%) missing values were detected across all samples. Since the percentage of missing values was low, it was assumed that the potential impact of missing values is insignificant [36]. Therefore, the remaining 1112 mass features were used for PCA (Figure 7A). To confirm the validity of this approach, PCA was also conducted on the dataset after missing value imputation using the K-nearest neighbor (KNN) method and showed similar indistinct grouping of the injected and control mice (Figure S8A).

In addition to PCA, unsupervised hierarchical clustering analysis (HCA) was conducted and a heatmap was generated to examine variations in metabolic patterns between the Aha and control groups (Figure 7B). HCA and heatmap visualization showed no clustering between the biological replicates of each group and no distinct differential abundance patterns between the two groups. This result further establishes that there are no substantial metabolic differences between control and injected mice. Similar to PCA, HCA performed using the dataset imputed via the KNN method resulted in indistinct clustering of the mice (Figure S8B).

Following global analysis using PCA and HCA, Student’s *t*-test was employed to identify metabolites that were differentially regulated between the two groups. In this analysis, a total of 15 out of 1112 metabolites were differentially abundant using a *p*-value of 0.05 and a fold change of ≥ 2 as cut-offs (Figure 7C). Of the 15 metabolites, 3 were upregulated and 12 were downregulated in Aha-treated mice compared to the control. Searching the 15 metabolites in the METLIN metabolite database using a mass tolerance of 5 ppm did not identify any known metabolite. The presence of a large number of unknown mass features is an intrinsic characteristic of untargeted metabolomic studies due to the complexity of the mammalian metabolome and the lack of structure characterization of a large number of metabolites (35, 36). Yet, the fact that only ~1.3% of metabolites were dysregulated, and that these dysregulated metabolites did not belong to any of the known major metabolic pathways, signifies that minimal metabolic alterations occur due to Aha administration.

Finally, to identify the metabolic pathways covered by the LC-MS analysis, analysis of equally-expressed metabolites was conducted using Metaboanalyst and the Kyoto Encyclopedia of Genes and Genomes (KEGG) metabolic pathway database (37). Several metabolic pathways were detected, including arachidonic acid metabolism, vitamin B6 metabolism, valine, leucine and isoleucine biosynthesis, galactose metabolism, and cysteine and Met metabolism (Table S9). This result indicates that the LC-MS analysis identified metabolites that belong to various metabolic pathways and that these pathways are not significantly changed in mice injected with Aha.

Collectively, the metabolomic analyses demonstrate that Aha administration does not significantly alter the murine plasma metabolome. This is in agreement with a recent study that investigated the metabolic effect of growing *E.coli* in media supplemented with ncAAs (38). A major advantage of our labeling technique is that it does not involve Met restriction or depletion as the case with other labeling strategies that use a Aha-enriched Met-free diet. Met dietary restriction has been shown to alter the metabolism in mouse models and in humans (39–41). Being a principle sulfur-containing amino acid, Met restriction specifically alters Met and sulfur metabolism (40, 41). Notably, the results of the pathway analysis identified several unchanged metabolites such as L-homocysteine, 5’-Methylthioadenosine, and 3-sulfinoalanine that belong to cysteine (another sulfur-containing amino acid) and Met metabolism (Table S9), indicating the advantage of the injection method with regards to its potential impact on metabolic functions.

## Methods

### Animal Model

Animals used in these studies were derived from female age-matched wild-type C57BL/6 mice (*Mus musculus*) purchased from The Jackson Laboratory. All experimental protocols were performed in compliance with established guidelines and all methods were approved by Purdue Animal Care and Use Committee (PACUC, protocols# 1209000723 and 1801001682). PACUC requires that all animal programs, procedures, and facilities at Purdue University abide by the policies, recommendations, guidelines, and regulations of the United States Department of Agriculture (USDA) and the United States Public Health Service (USPHS) in accordance with the Animal Welfare Act and Purdue’s Animal Welfare Assurance.

### Aha Injection, and Plasma and Tissue Collection

L-azidohomoalanine (Aha; Click Chemistry Tools) was resuspended in 1 × phosphate buffered saline (PBS) to 10 mg·mL^−1^, adjusted to pH 7.4, sterile filtered and stored at −20°C. All Aha injections were administered subcutaneously at 0.1 mg·g^−1^ total mouse weight. Mice (n = 3, biological replicates) were euthanized 0.5, 1, 2, 4, 6, 12 and 24 h post injection (hpi). Blood was harvested by cardiac puncture, collected in EDTA-treated tubes and centrifuged at 1,500 × *g* for 10 min at 4°C. The supernatant (plasma) was transferred into a new tube, snap frozen in liquid nitrogen and stored at −80°C. Liver, brain, kidney and hindlimb skeletal muscle tissues were dissected at each time point, snap frozen in liquid nitrogen and stored at −80°C. Control plasma and tissues were collected as described above from non-injected mice (n = 3 biological replicates). For the validation of model predictive ability, two Aha dosing regimens were used: (1) 12 h repeated doses (hrd) and (2) 24 hrd. Liver and brain tissues (n = 3 biological replicates) were dissected as described above at 6, 18, and 32 hpi, snap frozen in liquid nitrogen and stored at −80°C.

### Sample Preparation for Aha Analysis

For plasma sample preparation, 50 μL of plasma were mixed with 10 μL of 1 × PBS, pH 7.4, and 5 μL of 100 ng·μL^−1^ L-α-aminobutyric acid (α-ABA; Sigma Aldrich) that was used as an internal standard. 12.5 μL of trichloroacetic acid (TCA; Sigma Aldrich) were added to the mixture to precipitate proteins. The mixture was incubated for 10 min at 4°C and centrifuged at 16,000 × *g* for 10 min at RT. The supernatant was then mixed with 100% acetonitrile (ACN; Fisher Scientific) at a 1:1 ratio (v/v). The mixture was transferred to an HPLC autosampler vial for LC-MS/MS analysis. For calibration curve generation, Aha standards were prepared by mixing 50 μL of non-injected plasma with 10 μL of a known concentration of Aha and 5 μL of α-ABA. Proteins were then precipitated with TCA and prepared for LC-MS/MS analysis as described above.

For tissue sample preparation, tissues were rinsed with ice-cold 1 × PBS, pH 7.4 to remove residual blood and homogenized in ice-cold 1 × PBS, pH 7.4 using a TissueRuptor (Qiagen). The final homogenate weight was measured and converted to volume by using a homogenate density of 1 g·mL^−1^. Samples were then prepared for LC-MS/MS analysis as described for plasma by using 50 μL of the tissue homogenate. The remaining plasma samples and tissue homogenates were snap frozen and stored at −80°C until use for western blot and untargeted metabolomic analyses as described below.

### LC-MS/MS Targeted Analysis of Aha

An Agilent 1260 Rapid Resolution liquid chromatography (LC) system coupled to an Agilent 6470 series QQQ mass spectrometer was used for Aha analysis (Agilent Technologies). An Intrada Amino Acid 2.0 mm x 150 mm, 3.0 μm column (Imtakt Corporation) was used for LC separation. The buffers were (A) ACN, 0.3 % formic acid (FA; Sigma Aldrich) and (B) ACN/100 mM ammonium formate (20/80 v/v). The linear LC gradient was as follows: time 0 min, 20 % B; time 5 min, 20 % B; time 11 min, 35 % B; time 20 min, 100 % B; time 22 min, 100 % B; time 22.5 min, 20 % B; time 30 min, 20% B. The flow rate was 0.3 mL·min^−1^. Multiple reaction monitoring (MRM) was used for MS analysis. Data were acquired in a positive electrospray ionization (ESI) model based upon parameters in Table 3. The jet stream ESI interface had a gas temperature of 325°C, gas flow rate of 9 L·min^−1^, nebulizer pressure of 35 psi, sheath gas temperature of 250°C, sheath gas flow rate of 7 L·min^−1^, capillary voltage of 3500 V in a positive mode, and nozzle voltage of 1000 V. The delta electron multiplier voltage was 300 V. Agilent MassHunter Quantitative Analysis software was used for data analysis (v.8.0).

**Table 3.** Multiple reaction monitoring (MRM) table for amino acid LC-MS/MS data acquisition

| Compound name | Precursor ion (*m/z*) | Product ion (*m/z*) | Collision energy (eV) |
| --- | --- | --- | --- |
| Aha | 145.1 | 101.3 | 5 |
| Aha | 145.1 | 71.3 | 10 |
| Aha | 145.1 | 58.3 | 40 |
| Ala | 90 | 44 | 15 |
| Arg | 175 | 116 | 18 |
| Asn | 133 | 87 | 12 |
| Asp | 134 | 88 | 14 |
| Cys | 122 | 76 | 15 |
| Cys-Cys | 241.1 | 152 | 15 |
| Gln | 147 | 84 | 22 |
| Glu | 148 | 130 | 12 |
| Gly | 76 | 30 | 15 |
| His | 156 | 110 | 19 |
| Ile | 132 | 86 | 15 |
| Leu | 132 | 86 | 15 |
| Lys | 147 | 84 | 20 |
| Met | 150 | 104 | 15 |
| Phe | 166 | 120 | 15 |
| Pro | 116 | 70 | 15 |

### Western Blot Analysis of Aha-Labeled Tissues

Tissue homogenates were thawed and protein concentration was measured using the Pierce 660 nm Protein Assay (ThermoFisher Scientific). 200 μg of tissue homogenate was alkylated with 40 mM iodoacetamide for 30 min at RT in the dark with end-over-end rotation. Samples were then reacted for 2 h at RT with the following click reagents: 50 μM biotin-alkyne (ThermoFisher Scientific), 5 mM tris(3-hydroxypropyltriazolylmethyl)amine (THPTA; Click Chemistry Tools), 2 mM copper sulfate, 20 mM aminoguanidine and 10 mM sodium ascorbate. Following the click reaction, proteins were precipitated by adding ice-cold 100% acetone to the samples at a 4:1 ratio (v/v). Samples were incubated overnight at −20°C, centrifuged at 21,100 × g for 20 min at 4°C, supernatants were discarded, and protein pellets were vacuum-dried for 15 min at RT using a CentriVap (Labconco). Dried pellets were resuspended in 8 M urea in 1× PBS and centrifuged at 16,000 × g for 15 min at RT to remove insoluble particles. The supernatants were transferred into new tubes and protein concentration was measured using the Pierce 660 nm Protein Assay (ThermoFisher Scientific). Proteins were resolved on 4 – 20% SDS-PAGE gels (BioRad), transferred to a PVDF membrane (ThermoFisher Scientific) using the Trans-Blot Turbo Transfer System (BioRad) and probed overnight at 4°C with IRDye 680 Streptavidin (LICOR) diluted 1:3000 in 1:1 TBST:Blocking Buffer (BioRad). Membranes were imaged using an Azure Biosystems c600. Western blot images were analyzed using ImageJ (National Institutes of Health) to calculate the mean fluorescence intensities of each time point. The intensity of the control sample was used to normalize the intensity of each time point (n = 3 biological replicates per blot).

### Kinetic Modelling of Aha Distribution

Simulations were run on a Lenovo Yoga with an Intel Core i7-8550U CPU @ 1.8 GHz and 8 GB RAM. Simulations were performed using custom modeling scripts written in Python 3.6 (Supplement S1). Systems of ordinary differential equations (Supplement S2) were solved using a flexible high order solver from the SciPy python package (42). Most parameter values and ranges for fitting were informed from reported literature values or experimental measurements from this study. For parameters related to an experimental output ([fAha] or rF) without a reported literature value, an initial best estimate was selected to produce a single time-step change one order of magnitude lower than the maximum recorded experimental value. These parameters were then allowed to fit within a range of 1.5 orders of magnitude from the initial estimate. Parameters were fit with a least squares minimization algorithm from ‘Lmfit’, a prebuilt python library (43). All best fit values and boundary conditions can be found in the parameter tables (Supplements S3-5).

### Parameter Sensitivity Analysis and Model Validation

To effectively sample the input parameter space, Latin hypercube sampling (LHS) was utilized to generate unique parameter sets (n = 10000), sweeping each parameter value through a range defined by the boundary constraints from literature (Tables S3-S5) as previously detailed (32, 33). Global sensitivity analysis was performed on the LHS generated parameter sets using partial rank correlation coefficient (PRCC) analysis. This analysis quantifies the sensitivity of an output variable on the variation in input parameter values (32, 33). Here, PRCCs were determined for each of the 19 fitted parameters (Table S3) and 12 static parameters (Table S4) in the biodistribution model, as well as for all 8 parameters in the protein incorporation model (Table S5). PRCCs were used to characterize the influence of each parameter on the sum of square errors (SSE), the optimization metric for non-linear regression. Simulations used to inform PRCCs were performed on the Brown Supercomputing Community Cluster at Purdue University (44), with each simulation run on a single node with dual 12-core Intel Xeon Gold “Sky Lake” CPUs @ 2.60 GHz and 96 GB of memory.

The standard error of fitting was determined for the 19 fitted parameters (Table S3) in the biodistribution model and for all 8 fitted parameters (Table S5) in the protein incorporation model. Standard error values were determined from the covariance matrix during non-linear regression using the built-in functionalities of the ‘Lmfit’ python library (43).

### Plasma Sample Preparation for Untargeted Metabolomic Analysis

The plasma metabolome of non-injected control samples (n = 3 biological replicates) and samples collected 24 h post Aha injection (n = 3 biological replicates) was extracted by adding methanol: chloroform: water (1:1:1 v/v) to 80 μL of each plasma sample. Samples were vortexed briefly and centrifuged at 8,000 × *g* for 5 min at RT. The upper layer was transferred into a new tube and vacuum-dried overnight at RT. The dried fraction was reconstituted in 75 μL 5% ACN and 0.1% FA. Reconstituted samples were sonicated for 5 min, centrifuged at 16,000 × *g* for 8 min at RT, and the supernatants were transferred to HPLC autosampler vials.

### Untargeted LC-MS Metabolomic Analysis

Separations were performed on an Agilent 1290 UPLC system (Agilent Technologies). The metabolites were analyzed using a Waters Acquity HSS T3 column (1.8 μm, 2.1 × 100 mm), with a mobile phase flow rate of 0.45 mL·min^−1^, where the mobile phase A and B were 0.1% FA in double distilled water and ACN at a 1:1 ratio, respectively. Initial conditions were 100:0 A:B, held for 1 minute, followed by a linear gradient to 20:80 at 16 min, then 5:95 at 22.5 min. Column re-equilibration was performed by returning to 100:0 A:B at 23.5 min and holding until 28.5 min.

The mass analysis was obtained using an Agilent 6545 Quadrupole Time of Flight (Q-TOF) MS with ESI capillary voltage +3.2 kV, nitrogen gas temperature 325 °C, drying gas flow rate 8.0 L·min^−1^, nebulizer gas pressure 30 psig, fragmentor voltage 130 V, skimmer 45 V, and OCT RF 750 V. MS data scans (*m/z* 70-1000) were collected using Agilent MassHunter Acquisition software (v.B.06). Mass accuracy was improved by infusing Agilent Reference Mass Correction Solution (G1969-85001). MS/MS was performed in a data-dependent acquisition mode on composite samples.

### Metabolomic Data Statistical Analysis

Peak deconvolution and integration were performed using Agilent ProFinder (v.10.0). Bioinformatic analyses were performed using Agilent Mass Profiler Professional (v.13.1). Chromatographic peaks were aligned across all samples. Peak areas were normalized by log_2_-transformation and applying a 75% percentile shift. Metabolites were filtered out if present in only one sample. Furthermore, only metabolites present in all 3 replicates of either the control or Aha injected samples were included. Statistical analysis was performed using unpaired student’s *t*-test. Metabolites with *P* < 0.05 and fold change ≥ 2 were considered significant. Peak annotations were performed using the METLIN metabolite database, with a mass error of less than 5 ppm. Identifications were aided by MS/MS spectra comparisons. Principal component analysis (PCA), hierarchal clustering analysis (HCA) and metabolic pathway analysis were performed using MetaboAnalyst v.5.0.

## Conclusions

Here, we report for the first time the biodistribution kinetics of the widely used Met analog, Aha, in murine tissues, as well as the associated relative rates of incorporation of Aha into protein via protein synthesis and loss via metabolism and protein turnover. These results showed that liver and kidney have faster protein synthesis and turnover rates compared to brain and skeletal muscle, which is consistent with previous studies that utilized isotope labeling (29). We also demonstrated that subcutaneous injection allows for observing maximum protein labeling in a relatively short time (~ 6 h), which enables studying proteins with shorter half-lives, in contrast to the traditional method of introducing the ncAA in diet or using isotope-labeled amino acids. To support these findings, we developed a mathematical framework that described the distribution kinetics of Aha in murine tissues and its relation to the degree of protein labeling and computed the relative rates of protein synthesis and turnover. We further validated this framework for predictive modeling of Aha labeling against an experimental dataset including two different repeated injection dosing paradigms to demonstrate its efficacy as a tool for future experimental design. Finally, we investigated the impact of Aha administration on the plasma metabolome and demonstrated that Aha incorporation into cellular proteins does not have adverse effects on the normal physiology of mice. This observation further confirms previous results from our group that demonstrated that ncAAs do not affect the gross behavior nor the physical appearance of treated mice.

## Abbreviations

Aha: azidohomoalanine
HCA: hierarchical clustering analysis
hpi: hours post injection
KEGG: kyoto encyclopedia of genes and genomes
KNN: k-nearest neighbor
LC-MS/MS: liquid chromatography tandem-mass spectrometry
LHS: Latin hypercube sampling
Met: methionine
MRM: multiple reaction monitoring
ncAA: non-canonical amino acid
NSP: newly synthesized protein
ODE: ordinary differential equation
PCA: principal component analysis
PRCC: partial rank correlation coefficient
rF: relative fluorescence
SEf: standard error of fitting
fAha: free Aha
pAha: proteinous Aha
Sysrv: systemic venous reservoir

## Data Availability

All relevant data are within the manuscript and its Supporting Information files. Model and code files can be found at our lab’s GitHub repository (https://github.itap.purdue.edu/TamaraKinzerursemGroup/ncAABiokinetics)

## Acknowledgements

The authors thank Robyn McCain, Amber Jannasch and Bruce Cooper at Purdue Bindley Bioscience Center. The authors would also like to thank Karin F.K. Ejendal for care and maintenance of research mice. This work was supported by the National Institutes of Health [R01 AR071359 to SC and TKU] and National Science Foundation [Grant No. 1752366 to TKU]. The content is solely the responsibility of the authors and does not necessarily represent the official view of the NIH nor NSF.

## Supporting Information Captions

**S1 Text.** Code and model files.

**S2 Text.** Systems of ODEs for Aha model.

**S3 Table.** Distribution model fitting parameters.

**S4 Table.** Distribution model static parameters.

**S5 Table.** Incorporation model fitting parameters.

**S6 Figure.** Representative western blots informing Figures 2 and 3.

**S7 Figure.** Representative western blots informing Figure 6.

**S8 Figure.** Metabolomic analysis results following missing value imputation.

**S9 Table.** Selected metabolic pathways identified by Metaboanalyst analysis of the untargeted LC-MS analysis of Aha metabolome.

**S10 Table.** Untargeted LC-MS Aha metabolomic analysis raw data.

